# Discovery of a novel human antibody VH domain with potent activity against mesothelin expressing cancer cells in both CAR T-cell and antibody drug conjugate formats

**DOI:** 10.1101/2022.04.07.487497

**Authors:** Zehua Sun, Xiaojie Chu, Cynthia Adams, Tatiana V. Ilina, Michel Guerrero, Guowu Lin, Chuan Chen, Dontcho Jelev, Rieko Ishima, Wei Li, John W Mellors, Guillermo Calero, Dimiter S Dimitrov

**Author notes:** Corresponding authors. Dr. Zehua Sun,; Dr. Guillermo Calero,; Dimiter S Dimitrov. Authors contribute equally to the manuscript (Zehua Sun, Xiaojie Chu, Cynthia Adams).

## Abstract

Antibody based therapeutics targeting mesothelin (MSLN) have shown limited anticancer activity in clinical trials. Novel antibodies with high affinity and better therapeutic properties are needed. In the current study, we have isolated and characterized a novel VH domain 3C9 from a large size human immunoglobulin heavy chain variable (VH) domain library. 3C9 exhibited high affinity [KD (dissociation constant) < 3 nM] and binding specificity in a membrane proteome array (MPA). In a mouse xenograft model, 3C9 fused to human Fc became visible at tumor sites as early as 8 hours post infusion and persisted at the tumor site for more than 10 days. Both CAR-T cells and antibody domain drug conjugations (DDCs) generated with 3C9 were highly effective at killing MSLN positive cells in vitro without off-target effects. The X-ray crystal structure of full-length MSLN in complex with 3C9 reveals interaction of the 3C9 domains with two distinctive residues patches on the MSLN surface. 3C9 fused to human Fc domain drug conjugate was efficacious to inhibit tumor growth in a mouse xenograft model. This newly discovered VH antibody domain holds promise as a therapeutic candidate for MSLN-expressing cancers.

## INTRODUCTION

Mesothelin (MSLN) is an attractive tumor-associated antigen target for multiple solid tumors [1–4]. However, several antibody-based therapeutics including Immunotoxins, chimeric antigen receptor T-cells (CAR-Ts) and antibody drug conjugates (ADCs) have shown limited anti-tumor activity in clinical trials [1, 5, 6], indicating the need for new therapeutic approaches including more specific anti-MSLN antibodies. There are several unmet challenges for antibody-based therapeutics against MSLN-expressing solid tumors: (1) higher antibody affinity and avidity allowing recognition of tumor cells with low surface density of MSLN; (2) improved antigen specificity to minimize off-target binding and cytotoxicity; (3) better antibody penetration of solid tumors; and (4) less aggregation of the antibody.

Since the first antibody domain caplacizumab was approved by the FDA in 2019, VH antibody domains have been widely discussed for their multiple potential advantages over traditional antibodies, especially better penetration of solid tumors because of small size and reduced immunogenicity (fewer epitopes) [7–10]. In theory, antibody domains, which are about 10 times smaller than full-size IgGs, could have up to a 100-fold-higher effective diffusion coefficient into solid tumors [7, 10]. However, their half-life in circulation is shorter than that of full-size antibodies and they can be cleared quickly if binding to the target is not of high affinity. Discovery of novel antibody domains with high affinity and engineering potential to prolong half-life could be beneficial for the development of therapeutics directed against MSLN. To date, antibody domains to MSLN have not been reported to enter clinical development.

Previously, we constructed a large-scale human antibody VH domain library based on thermo-stable anti-aggregation scaffolds for phage display [11, 12]. Panels of binders were isolated from this antibody VH domain library against different cancer targets, demonstrating the quality and diversity of the library. In the present study, we describe the isolation and characterization of a novel MSLN specific antibody domain - 3C9 – with high affinity [KD (dissociation constant) < 3 nM] and no off-target binding detected in a human membrane proteome array of ~6,000 surface proteins. The crystal structure of full-length MSLN bound to 3C9 complex revealed a unique binding site of 3C9 to MSLN, which differs from other MSLN binders including those that have advanced to clinical development. ADC and CAR-T cells containing 3C9 as the binding motif show potent cytotoxicity against MSLN-expressing tumor cells without off-target effects.

## RESULTS

### Discovery and characterization of a novel human MSLN-specific VH antibody domain

We designed human recombinant MSLN constructs, which were expressed and used as antigens to isolate MSLN-specific binders from a large size human antibody VH domain library by phage display [12]. This approach yielded a panel of unique domains with the affinities ranging from 0.5 nM to 200 nM for human MSLN. We extensively characterized one VH domain, 3C9, with high affinity and favorable non-aggregation profile ([12], and Figure 1A). In an ELISA assay, 3C9 exhibited an EC50 less than 10 nM (Figure 1B). In a Surface Plasmon Resonance (SPR) assay, 3C9 exhibited a high affinity to human MSLN with the KD less than 3 nM (Figure 1E). VH 3C9 also binds macaque MSLN, though with a lower binding affinity (KD > 100 nM, data not shown).

**Figure 1.**
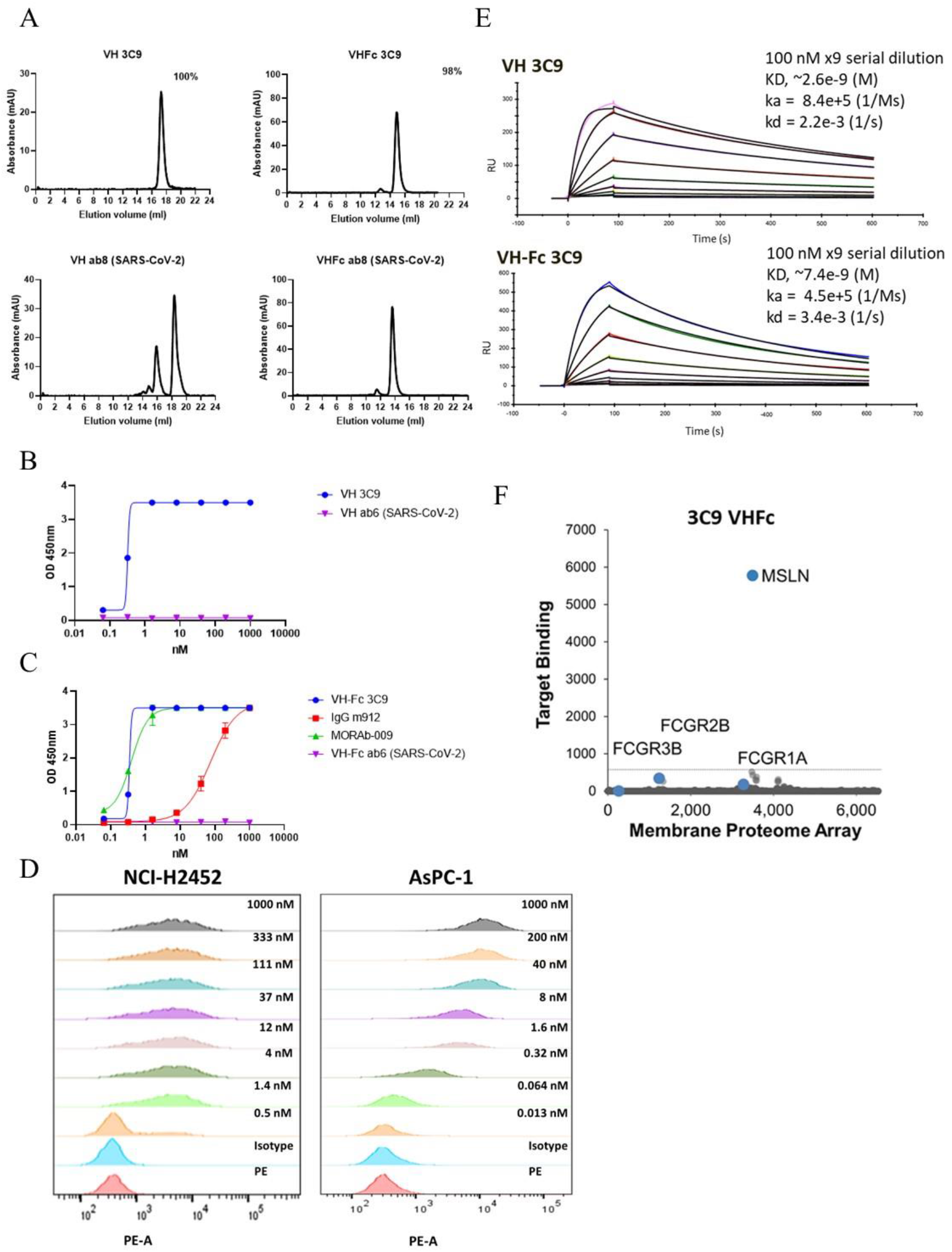
Characterization of 3C9. A. SEC data of both VH and VH-Fc 3C9. Aggregation resistant VH and VH-Fc ab8 were used as controls. B-C. ELISA of VH/VH-Fc binders binding to human MSLN. D. Cell specific binding assay by VH-Fc 3C9 in a dose dependent manner. E. SPR of 3C9 VH and the VH-Fc forms. F. Lack of non-specific binding measured by a Membrane Proteome Array (MPA). Antibody domain 3C9 was fused human Fc protein for the examination by flow cytometry. VH-Fc 3C9 (20 μg/ml) was tested in a Membrane Proteome Array against 6,000 different human membrane proteins.

To measure the effect of avidity and extend the half-life, the VH domain was fused with human IgG1 Fc protein to form proteins of a bivalent format (VH-Fc). VH-Fc proteins were purified from transfected Expi293 cells with yields ranging 40-60 mg/L. VH-Fc 3C9 has an EC50 of less than 1 nM (Figure 1C) by ELISA, which was comparable to amatuximab (MORAb-009), a mouse-human IgG1/k monoclonal antibody in Phase 2 clinical development[13]. By SPR assay, the KD of VH-Fc 3C9 was 7 nM (Figure 1E). The reduced avidity is probably caused by the fixed orientations of VHs by the Fc protein. VH-Fc 3C9 specifically binds MSLN positive cell lines in a concentration dependent manner (Figure 1D).

A membrane proteome array (MPA) platform [8]was used to test specificities of VH-Fc 3C9 against a total 6,000 different human membrane proteins including 94% of all single-pass, multi-pass, and GPI-anchored proteins (GPCRs, ion channels, and transporters). Cell specific binding was first verified in a concentration dependent manner and was validated before the MPA assay. VH-Fc 3C9 showed no off-target binding to 6,000 membrane associated proteins tested (Figure 1F).

### Structural basis of 3C9 binding to MSLN

Crystals of the SEC purified MSLN and 3C9 complex appeared under 0.18 M tri-ammonium citrate and 20% PEG 3350. Initial phases to 2.9 Å resolution were obtained by multiple anomalous diffraction (MAD) of selenomethionyl labeled VH 3C9. At this resolution, side chains were clearly visible, allowing residue placement, as described below. (The model was subjected to several rounds of manual building in Coot64 and refinement using Phenix66 to a final R-free and R-work of 25 and 22%, respectively (Table 1). The asymmetric unit contains two mesothelin molecules (Figure S1A), each one interacting with two VH 3C9 molecules (Figure 2A and Figure S1B). The structure of the full mature MSLN (residues 300 to 592) comprises 10 alpha helical hairpins (helix-loop-helix) forming a super-helical solenoid (Figure S2A). An internal disulfide bond, C442-C468, (Figure S2A) between hairpins 5 and 6 confers rigidity to the solenoid. Calculation of the surface electrostatic potential shows the presence of charged pockets and crevices that could represent prospective interaction areas (Figure S2B).

**Table 1.** Data collection, phasing and refinement statistics.

|  | SSRL (12-1) |
| --- | --- |
| **Data collection** |  |
| Space group | P4<sub>3</sub>22 |
| Cell dimensions |  |
| *a*, *b*, *c* (Å) | 114.1 114.1 340.7 |
| $\alpha$, $\beta$, $\gamma$ (°) | 90, 90, 90 |
| Resolution (Å) | 60-2.9 |
| *R*<sub>sym</sub> or *R*<sub>merge</sub> | 0.008(0.45) |
| *I* / $\sigma I$ – CC1/2 | 27.5(1.1) – 99(35) |
| Completeness (%) | 100(100) |
| Redundancy | 25(18) |
| **Phasing** |  |
| Wavelength (Se peak) | 0.9794 |
| No of sites | 8 |
| FOM (SAD) | 0.27 |
| **Refinement** |  |
| Resolution (Å) | 37-2.9 |
| No. reflections | 51008 |
| *R*<sub>work</sub> / *R*<sub>free</sub> | 22.1/24.4 |
| *B*-factors |  |
| Protein | 65.5 |
| Ligand/ion | N/A |
| Water | 74 |
| R.m.s. deviations |  |
| Bond lengths (Å) | 0.009 |
| Bond angles (°) | 1.33 |

\*Values in parentheses are for highest-resolution shell.

**Figure 2.**
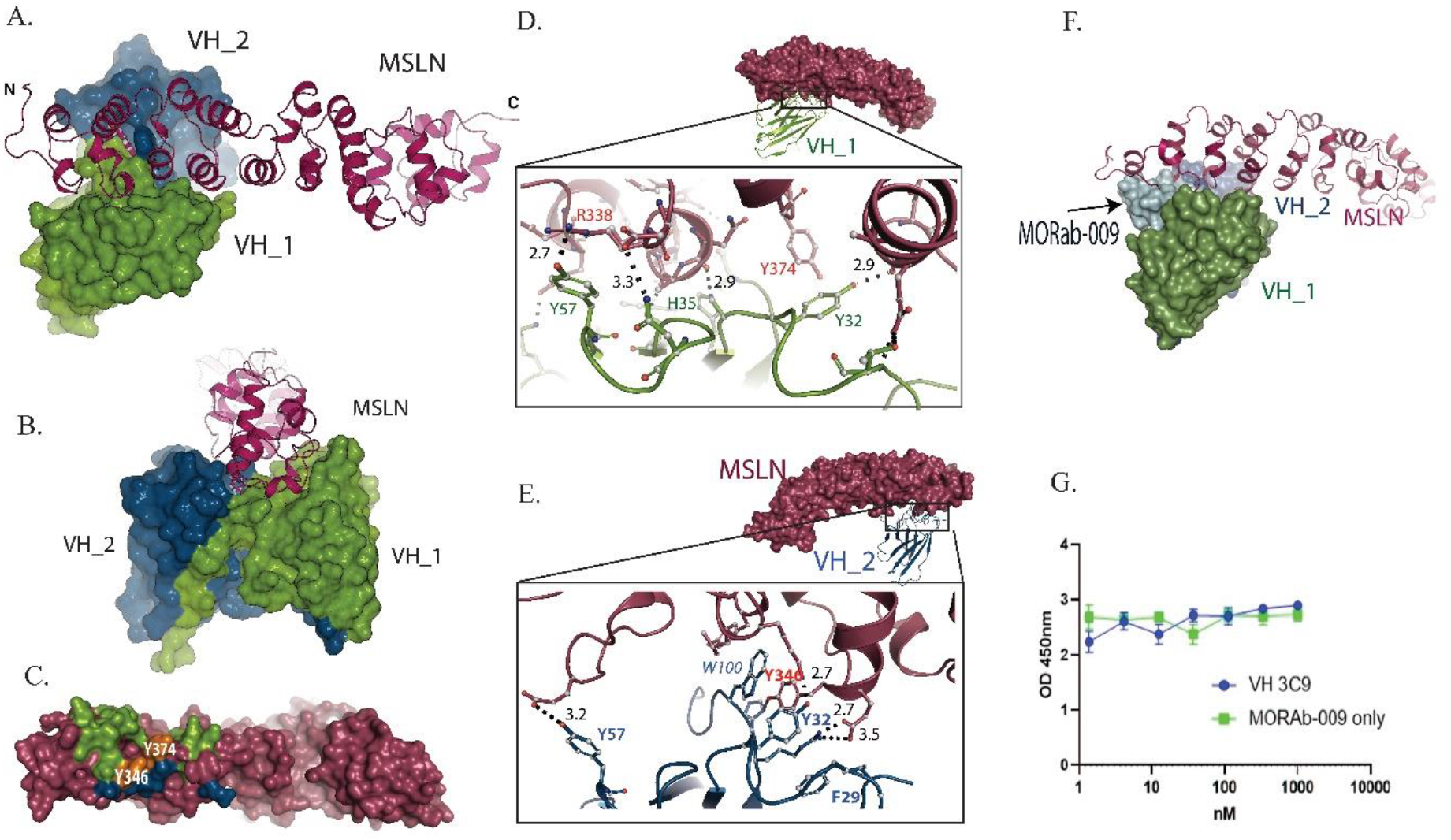
Structure of MSLN-3C9 complex, determined by crystallography. A. MSLN, (coordinate shown from residue 300 to 582, purple), complex with two VH 3C9 domains, VH_1 and VH_2 (green and blue, respectively). B. The same structure, from different orientation, highlighting swapping of the C-terminal beta-strands in two VH 3C9 domains. C. Map of VH_1 (green) and VH_2 (blue) interaction site on MSLN (purple) with highlight of the locations of Y346 and Y374 (orange). D. Side chain interaction of MSLN with VH_1 (green) with highlight of Y374 (orange). E. Side chain interaction of MSLN with VH_2 (blue) with highlight of Y346 (orange). F. Binding surface of MORab-009 (ice blue) and VH_1 and VH_2 interaction with MSLN, showing that MORAb-009 (Amatuximab) interacts more N-terminal region than VH 3C9. G. VH 3C9 and Amatuximab have no competition in competing ELISA. Expanded figures for panel A are shown in Figures S1 and S2, extended figures for panel B and C are shown in Figure S3, extended figures for panel D and E are shown in Figure S4, extended figures for panel F are shown in Figure S6, respectively.

Two 3C9 molecules (heretofore VH_1 and VH_2) form a swapped dimer domain that features a significant contact area (burying 1805 A2) and 31 hydrogen bonds/salt bridges (Figure 2B, 2C, S3A, and S3B). These interactions comprise the CDR3 loops (residues 97-104); and the C-terminal strand of VH_1 (VH_2) completing the B-sheet of VH_2 (VH_1) (Table S1, interactions and Figure S3C and S3D).

The N-terminal region of MSLN interacts with two VH 3C9 molecules, VH_1 and VH_2, burying a surface area of 799 A2 and 391 A2, respectively. MSLN interaction with the VH_1 domain involves hairpin loops 1-4 that contact CDR1 (residues 26-33), CDR2 (residues 51-57) and CDR3 loops, and residues from strands 3-5 (of the core B-sheet) that are contiguous to the CDR2 and CDR3 loops (Figure 2D, S4A-S4E, Table S1). Interactions with the VH_2 domain involve contacts between hairpin loops 2-4 and the CDR3 domain where W100 is buried in a small hydrophobic pocket formed by residues Y346, L349, Y374 and L377; and Y32 (CDR1 loop) forming a salt bridge with K378 (Figure 2E, S4F-S4H, and Table S1). The MSLN-3C9 interaction was validated by mutagenesis experiments; MSLN Y346A and Y374A mutants were tested since they have multiple interactions with both VH_1 and VH_2 (Figure 2C). The Y346A mutant and the double mutant Y346A and Y374A abolish 3C9 binding, while the Y374A mutant had a partial effect (Figure S5).

Overlay between the MSLN-3C9 (this work), and the MSLN N-terminal region (residues 303 to 359) with amatuximab Fab [14] structures shows that 3C9 and amatuximab MSLN interactions involve largely non-overlapping regions (Figure 2F and S6). Indeed, competitive ELISA assays show no MSLN binding competition between VH 3C9 and amatuximab (Figure 2G). The total buried surface area (1200 vs 950 A2) and the overall number of hydrogen bonds/salt bridges (21 vs 9) is larger for the MSLN-3C9 complex, explaining the enhanced affinity of 3C9 over amatuximab (Table S1).

### Pharmacokinetics and tumor delivery of VH-Fc 3C9

Tissue distribution and persistence time are important considerations in evaluating the therapeutic potential of an antibody. To investigate these properties, the in vivo pharmacokinetics of human IgG1 Fc fused 3C9 was evaluated in a mouse subcutaneous xenograft cancer model using AsPC-1-luciferase cells. 4-6 weeks nude mice were injected via s.c. with 5.0 × 10^5 AsPC-1-luciferase cells. Seven days later, tumors were visible and approximately 200 mm3 in size. 200 μg (~10 mg/kg, mice) VH-Fc 3C9 labeled with far infrared dye YF®750 SE (named as Ab-YF750) was injected i.p. into five mice. Images were recorded at 1h, 4h, 8h, 1d, 3d, 5d, 7d, 10d and 14d (Figure 1A). Infrared fluorescence in the tumor was detected as early as 8 hours after i.p. antibody administration. The VH-Fc 3C9 signal persisted in tumors for more than 10 days (Figure 3B-C). To confirm 3C9 enrichment, tumors were dissected at 3 and 14 days post antibody infusion (experiment endpoint) and assay for fluorescence, As shown in Figure 3B, the dissected tumor showed fluorescence signal of antibody enrichment in the tumor site.

**Figure 3.**
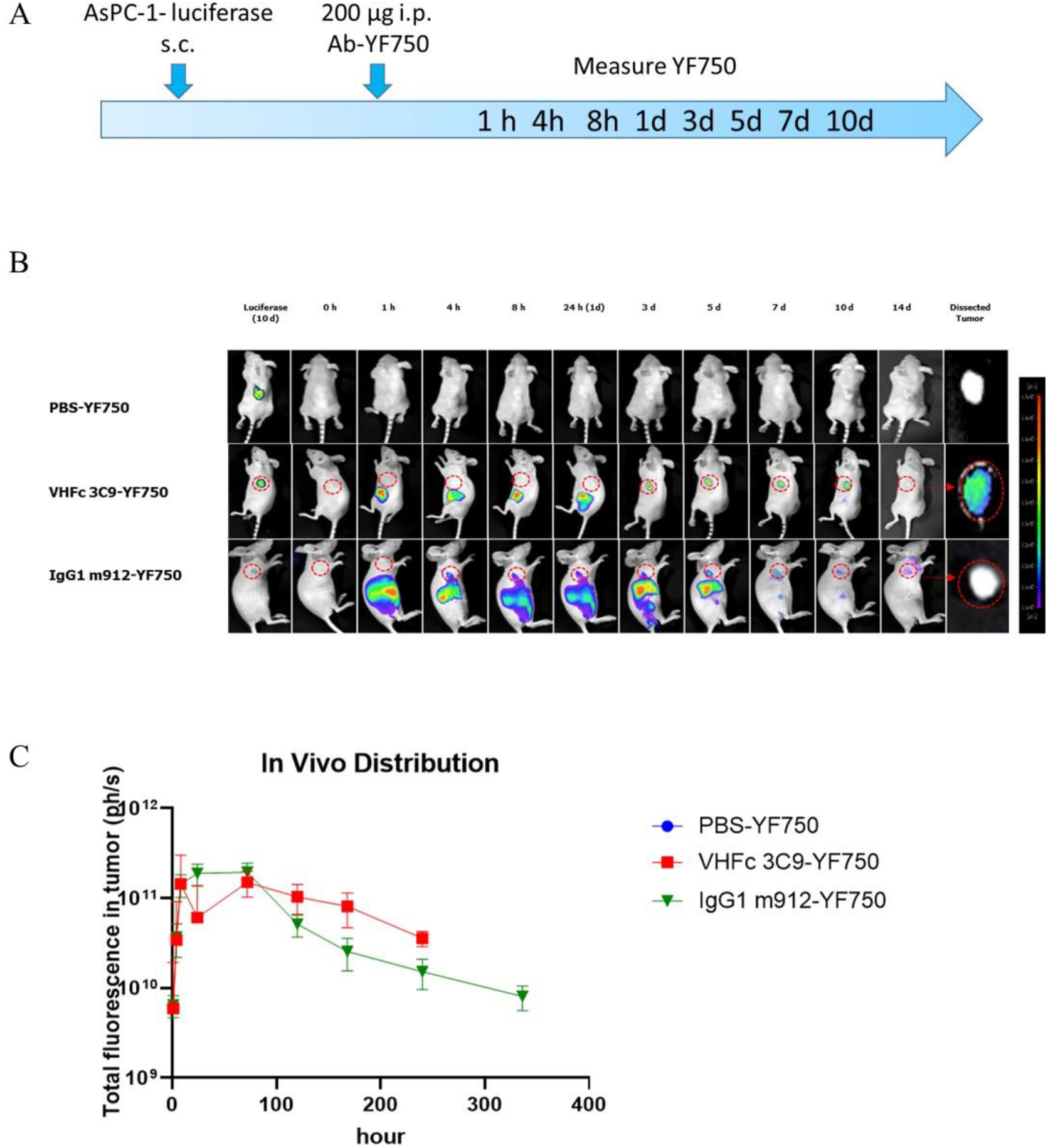
Pharmacokinetics of VH-Fc 3C9-YF750 by i.p. administration at the indicated time points. (A) Schematic view of the mice injection and data collection. (B) PBS-YF750 group, VH-Fc 3C9-YF750 group and IgG1 m912-YF750 group; Nude mice (n=5) were administrated with VH-Fc 3C9-YF750 or PBS-YF750 by the i.p. route. The fluorescence intensity at the tumor location as shown in the yellow dashed line circle of mice in (B) was measured at the indicated time point. To confirm and pinpoint the location of the tumor cells, the substrate was injected via i.p. at 10d to measure the signal of luciferase from AsPC-1-luciferase cells. To cofirm the antibody enriched in the tumor, the tumor was dissected at 14 dpi for YF750 measurement. (C) Total fluorescence in tumor (ph/s).Data are represented as mean ± SEM

### VH domain drug conjugates (DDC) exhibit tumor cytotoxicity effects both in vitro and in vivo

We previously reported that CAR-T with a MSLN-specific ScFv (m912) has therapeutic potential in both in vitro and in vivo studies [5]. To test the efficacy of CAR-T cells generated with VH domains, we constructed second generation CAR constructs based on the VH domain 3C9. Transduced T cells displayed ~ 60% CAR surface expression (Figure 4A). 3C9-CAR transduced T cells mediated antigen-specific killing of MSLN positive AsPC-1 and NCI-H2452 cells compared with untransduced T cells and did not kill MSLN negative 293T cells.

**Figure 4.**
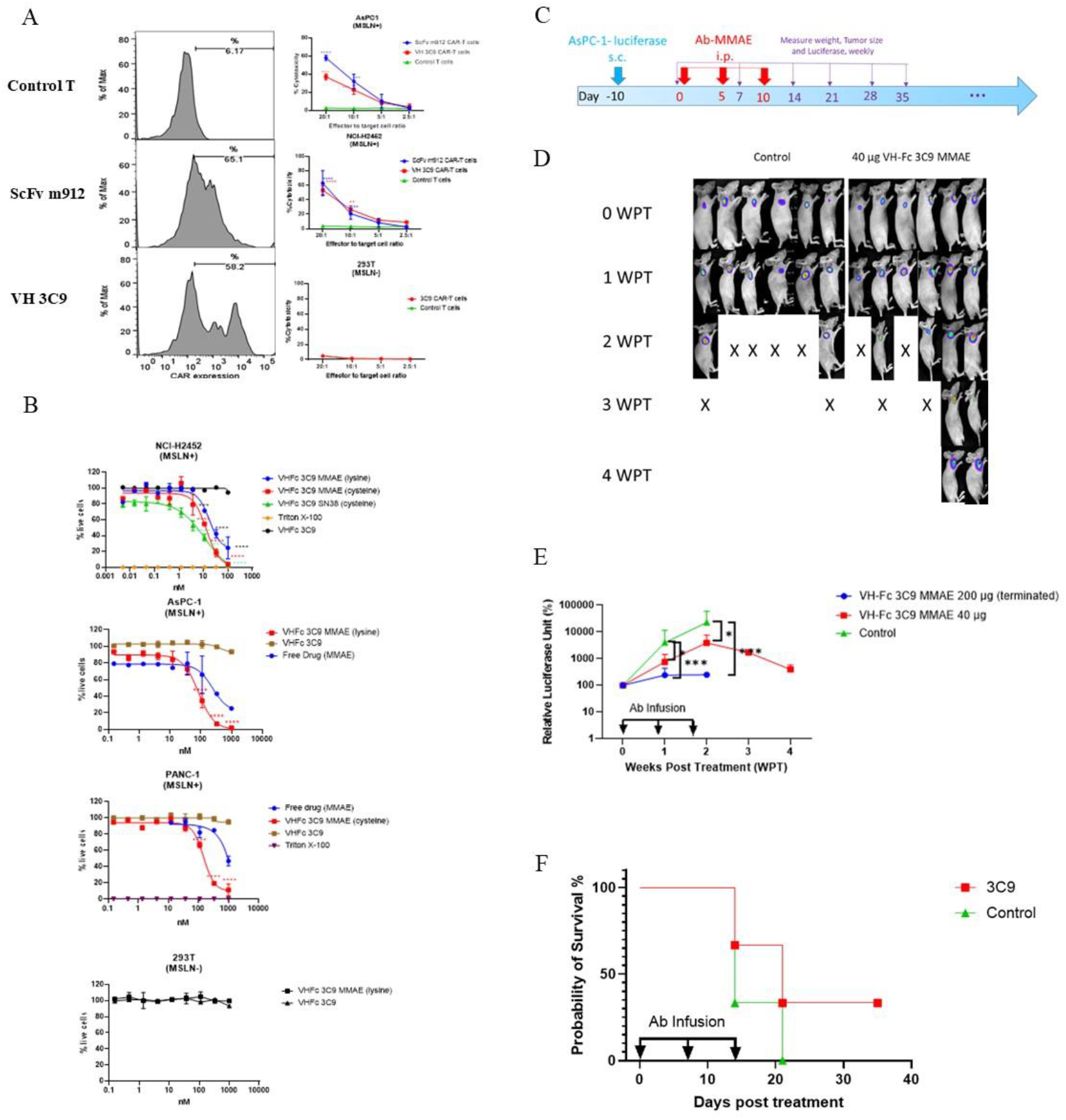
Characterization of MSLN-directed CAR T cells and DDCs *in vitro* toward MSLN positive human pancreatic cancer cell line and lung mesothelioma. A. VH domain/ScFv CARs expression measured by flow cytometry in CAR T cells three days after transduction. Percentage of lysis of MSLN+ tumor cell lines (NCI-H2452 and Aspc-1) by 3C9 CAR T cells measured by LDH release assays (n = 4–9). **, P < 0.01; ***, P<0.001; ****, P<0.0001. B. Efficacy of DDCs in killing tumor cells and non MSLN cells *in vitro*. C. Schematic view of the mice injection and data collection. D. Fluorescence measurements of control group and ADC group. E. Relative luciferase signals of tumor in control group and ADC groups. F. Survival rate measurement.

Domain drug conjugates to cytotoxic payloads (DCC) have been evaluated recently in different tumor models [12]. Exposed lysine-based conjugation is a widely used non-specific conjugation strategy for antibody drug conjugates with two lysine-based ADCs having been approved by the FDA [15–17]. Payload SN38 has proven to be highly effective in the ADC format with low off-target toxicity [18]. We expressed VH-Fc format antibody proteins for lysine conjugation with a protease-labile valine-citrulline linker (OSu-Glu-vc-PAB-) linked with monomethyl auristatin E (MMAE), and cysteine conjugation by a pH cleavable linker (CL2A) with SN-38 as payload to constitute VH-Fc 3C9 antibody DDCs. Both VH-Fc 3C9-MMAE and VH-Fc 3C9-SN-38 killed MSLN positive tumor cells \ without off-target effect in non-MSLN cells (Figure 4B).

We evaluated the antitumor activity of VH-Fc 3C9 MMAE (lysine conjugated, CAR is ~3.6) in vivo in an xenografted model of pancreatic cancer. In the xenogeneic tumor model of AsPC-1-Luc cells, VH-Fc 3C9 MMAE showed dose-dependent efficacy, and the highest dose for the studies, 10 mg/kg based on the ADC did affect mouse body weight, even though significant inhibition (p<0.0001) of tumor growth was observed. High DAR (~3.6) could be one possible reason causing toxic to administrated mice; thus, we terminated data collection of this high dose group. Treatment with 2 mg/kg VH-Fc 3C9 MMAE also resulted in significant inhibition of tumor growth (Figure 4C-D). At a dose of 2 mg/kg VH-Fc 3C9 MMAE, treatment of AsPC-1-Luc tumors showed a 33% survival rate, in comparison with 0% in control group (Figure 4E-F).

## DISCUSSION

Antibody-based therapeutics targeting MSLN-expressing tumors have shown limited activity in clinical trials. Amatuximab that has potent ADCC activity showed moderate antitumor activity (NCT02357147). Immunotoxins such as SS1P (NCT01362790) can elicit neutralizing antibodies which largely affects the efficacy. Promising antitumor activity of MSLN-specific CAR T-cell therapy in combination with anti-PD1 therapy was observed in malignant pleural mesothelioma (MPM) (NCT02414269). Moreover, a panel of antibody drug conjugates was also reported exhibiting manageable safety and encouraging preliminary antitumor activity in MSLN positive solid tumors (NCT01439152, NCT01469793 and NCT02341625).

Antibodies with satisfactory specificity, high affinity, and in vivo persistence are critical to successful therapeutic development. Specificity of mAbs is one key issue that impacts the efficacy of antibody drugs because non-specific interactions can lead to off-target binding which results in toxicity or fast antibody clearance in vivo [19]. 3C9 reported here did not bind the 6,000 human membrane-associated proteins in the MPA assay, indicating low potential for off-target toxicity. Although a potential limitation of human antibody VH domains is the lack of sufficient binding affinity required by therapeutic application, both VH and VH-Fc 3C9 showed less than a 10 nM dissociation constant against MSLN. Based on the structural study of binding site, larger binding surface of 3C9, compared to amatuximab Fab, explains the high binding affinity of VH 3C9. In addition, VH-Fc 3C9 accumulated at in vivo tumor sites as early as 8 hours after intraperitoneal administration and persisted in tumors for more than 10 days. To our knowledge, it is the first report of measurement of the pharmacokinetics of an antibody domain in vivo in a tumor bearing mice model. It is also the first time to report the analysis of full length MSLN based on the crystal of MSLN-antibody complex. The PDB file will be subject to submission after a peer reviewed journal accepts our manuscript.

The antitumor activity for anti-MSLN ADCs with low off-target toxicity is encouraging in recent studies. DMOT4039A (h7D9.v3) is an MMAE conjugated humanized IgG1 anti-mesothelin mAb with promising antitumor activity and an acceptable safety profile based on a completed phase I study (NCT01469793). Anetumab ravtansine, another anti-MSLN ADC linked to maytansinoid DM4 exhibited a manageable safety profile in heavily pretreated patients with mesothelin-expressing solid tumors [20]. Compared with full length IgGs, VH and VH-Fc antibody formats have a reduced size that can be beneficial for penetration of solid tumors. In the present study, MMAE or SN38 conjugated VH-Fc proteins (either by lysine-based conjugation or cysteine-based conjugation, respectively) exhibit specific killing of MSLN positive tumor cells without off-target killing of MSLN negative cells. The overexpression of mesothelin in pancreatic cancer was confirmed by detailed immunohistochemical studies ([21]). VH-Fc 3C9 MMAE inhibited tumor growth at a dose of 2mg/kg. Continuous optimization of VH-Fc 3C9 and conjugation with new drug have promise in new drug development against pancreatic cancer.

Several bi-paratopic antibodies have been reported in different tumor models [22–25]. In a breast cancer model, a bi-paratopic antibody shows enhanced avidity and cross-linking activity to promote HER2 clustering and lysosomal degradation [22]. Although the epitope of 3C9 is quite close to the epitopes of amatuximab Fab, and there is no competition observed between 3C9 and amatuximab, thus a biparatopic antibody including VH 3C9 and Fab amatuximab may further increase the avidity without significant increase of antibody size [26, 27].

Clinical trials using MSLN-directed CAR T cells using a single-chain variable fragment (ScFv) format have demonstrated potential but with limited effects [26]. No MSLN-directed CAR T cells based on the VH domain have been reported yet. We designed CAR constructs based on VH domain 3C9 recruiting 4-1BB as the signaling component. Although CD28 containing CAR-T cells show rapid elimination of tumors, 4-1BB containing CAR-T cells induce long-term remission with greater persistence [28]. In prostate cancer, 4–1BB-containing PSCA-directed CAR T cells show more durable antitumor responses compared with CD28-containing CAR T cells [29]. In ovarian cancer, both CD28- and 4–1BB-containing CAR T cells were able to control SKOV3 cells and prolong the mice survival [20]. In our study, 4–1BB-containing VH 3C9 CAR T cells were effective in inducing tumor toxicity in vitro with 40%-80% killing of both AsPC-1 cells (pancreatic cancer) and NCI-H2452 (mesothelioma) without off target toxicity, demonstrating the possibility of using a VH domain in a CAR construct.

Immune escape by tumors can occur through multiple mechanisms such as antigen loss or binding escaping mutations within the same antigen [24]. Multispecific CAR-T cells potentntly eliminate antigen-heterogeneous B cell tumors in preclinical models to conquer the relapse caused by antigen loss or down-regulation [23]. Bi-paratropic CARs or ADCs can target mutant variants simultaneously. Optimization of the CAR construct can further increase the efficacy. In one recent study, a CD19 specific CAR was designed inside T cell receptor–CD3 complex targeting cell surface antigens of low abundance (<20 antigens per cell) [24]. We are constructing 3C9 CAR-Ts using a similar strategy. Combining CAR therapy with the use of checkpoint blockades to overcome CAR T-cell inhibition is another option to increase the efficacy. One phase II study combining MSLN CAR (m912 ScFv based) T-cell treatment in MPM with pembroluzimab is ongoing (NCT02414269).

In summary, we isolated and characterized a novel antibody VH domain – 3C9 - with high affinity to MSLN, aggregation resistance, and high specificity. The 3C9 VH domain exhibited specific tumor cell killing in both CAR-T and domain drug conjugate (DDC) formats. Pharmacokinetic studies and animal trials of mono-topic/bi-paratopic CAR-T and ADCs are in progress to further evaluate clinical developability.

## MATERIALS AND METHODS

### Expression and purification of MSLN protein, VH binders, and VH bivalent proteins

The gene of human mesothelin was synthesized by IDT (Coralville, Iowa) with the sequence obtained from Uniprot (https://www.uniprot.org/uniprot/Q13421). The MSLN domain (residues 296-606) were cloned into an expression plasmid. This plasmid contains a CMV promotor and woodchuck posttranscriptional regulatory elements with a His tag. Proteins were purified by Ni-NTA (GE Healthcare). MSLN specific VH domains were in the pComb3x vector and purified from Escherichia coli HB2151 bacterial culture at 30°C for 16 h with stimulation by 1 mM IPTG. Cells were lysed by Polymyxin B (Sigma-Aldrich). Lysates were spun down and supernatant was loaded over Ni-NTA (GE Healthcare). For conversion to Fc-fusion, the VH gene was re-amplified and re-cloned into pSectaq vector containing human Fc. VH-Fc proteins were expressed in the Expi293 expression system (Thermo Fisher Scientific) and purified with protein A resin (GenScript). Buffer replacement in protein purification used Column PD 10 desalting column (GE Healthcare). All protein purity was estimated as >95% by SDS-PAGE and protein concentration was measured spectrophotometrically (NanoVue, GE Healthcare). Further details can be found in our previous publication [14].

### ELISA and SPR

For ELISA assays, antigen protein was coated on a 96-well plate (Costar) at 50 ng/well in PBS overnight at 4°C. For the soluble VH binding assay, horseradish peroxidase (HRP)-conjugated mouse anti-FLAG tag antibody (A8592, Sigma-Aldrich) was used to detect VH binding. For detection of human Fc protein, HRP-goat anti-human IgG Fc secondary antibody (A18817, Thermo Fisher Scientific) was used. For the competition ELISA, 200 nM of IgG1 m912, or 50 nM VH-Fc 3C9 was incubated with serially diluted VH proteins, and the mixtures were added to antigen-coated wells. After washing, competition was detected by HRP-goat anti-human IgG Fc secondary antibody (A18817, Thermo Fisher Scientific). The kinetics of the antibody fragments were determined using a Biacore X100 (GE Healthcare). Human MSLN (Advance BioMatrix 5123-0.1mg) was (10 mg/mL) was immobilized onto a CM5 sensor chip (GE Healthcare, BR100012) by amine coupling. The antibody fragments diluted in HBS-EP buffer (10 mM HEPES, 150 mM NaCl, 3 mM EDTA, and 0.005% surfactant P20, pH 7.4) were injected over an immobilized surface (200 - 400 RU) for 90 sec at a rate of 50 μL/min, followed by dissociation for 600 sec. After each sample injection, the surface was regenerated by injection of regeneration solution (10 mM Glycine/HCl buffer, with 10% Glycerol at pH 2.0 for VH and VH-Fc binders). The kinetic values, ka, kd, and KD were calculated using the BiacoreX100 Evaluation Software (GE Healthcare).

### ADC generation

Monomethyl auristatin E (MMAE) was conjugated to VH-Fc 3C9 via the cross-linker OSu-Glu-VC-PAB (SET0100, Levena Biopharma, USA) with a molar ratio mAb:OSu-Glu-VC-PAB-MMAE of 1:10. The conjugation was performed in buffer composed of 50 mM potassium phosphate, 50 mM sodium chloride, and 2 mM EDTA (pH 6.5) with the reaction run for 18-24 h at room temperature. The reaction was stopped by adding 50 mM sodium succinate (pH 5.0) followed by buffer replacement by using Column PD 10 desalting column (GE Healthcare). Antibody SN-38 drug conjugates were prepared using purified antibody [30]. Briefly, antibodies were reduced using a 100X molar excess of TCEP (Millipore-Sigma CAT#580561) followed by reformation using 20 equivalents of dehydroascorbic acid (Millipore Sigma CAT# 261556). Rather than adjusting pH using titration, buffers were changed using a PD-Minitrap G25 column (Cytiva CAT#28918007). Antibody was similarly diluted 1:1 (v/v) with propylene glycol (Millipore Sigma CAT# P4347) and 3 equivalents of CL2A-SN-38 (Cayman Chemical CAT#33941) were added and incubated overnight at 4°C. The protein was washed and concentrated using 30 kDa Amicon centrifugal filter unit (Millipore Sigma CAT#UFC8030).

### CAR-T cell preparation

M SLN-directed CAR constructs containing different human MSLN-specific VH followed by 4–1BB-CD3ζ signaling domain. T cells from healthy donors were transduced with γ-retroviral vectors encoding CAR constructs. CAR transduction efficiency was determined by flow cytometry with CAR expression detected by human MSLN staining. Retroviral supernatant production and activation and viral transduction of T cells was performed as described previously [25].

### Cell Viability Assays

Cell viability was measured using CellTiter-Glo or LDH-Glo (G7570, J2380, Promega). Briefly, MSLN positive or negative cells were plated into 96-wells, allowing attachment and growth for 24 hr, then triplicate wells were treated with ADCs, naked antibodies, free drugs, or ADCs plus competitor antibodies. Three to five days later, when untreated control wells were 70 to 90% confluent, reagent was added to the plates according to the supplier’s instructions. Wells treated identically but wells without cells were used to subtract background. Fluorescence (ex: 570 nm, Em: 585 nm) was measured using a CLARIOstar microplate reader (BMG Labtech) and data analyzed using GraphPad Prism 8 software. Significance was tested using one-way ANOVA, followed by the Tukey’s multiple post hoc test. ****P < 0.0001; ***P < 0.001; **P < 0.01; *P < 0.05 between the indicated groups. PD-L1 neutralization was done in a PD-1/PD-L1 Blockade Bioassay (J1250, Promega).

### Dynamic Light Scattering (DLS)

VH domains were buffer-changed to DPBS and filtered through a 0.22 μm filter. The concentration was adjusted to 5 mg/mL; 500 μL samples were incubated at 37°C. DLS measurement was performed on Zetasizer Nano ZS ZEN3600 (Malvern Instruments Limited, Westborough, MA) to determine the size distributions of protein particles.

### Size Exclusion Chromatography (SEC)

The Superdex 200 Increase 10/300 GL chromatography (GE Healthcare, Cat. No. 28990944) was used for loading samples. The column was calibrated with protein molecular mass standards of Ferritin (Mr 440 000 kDa), Aldolase (Mr 158 000 kDa), Conalbumin (Mr 75 000 kDa), Ovalbumin (Mr 44 000 kDa), Carbonic anhydrase (Mr 29 000 kDa), Ribonuclease A (Mr 13 700 kDa). 150 μl filtered proteins (1-2 mg/ml) in PBS were used for analysis. Flow rate was 0.4 ml/min.

### Membrane Proteome Array

Integral Molecular, Inc. (Philadelphia, PA) performed specificity testing of VH-Fc 3C9 using the Membrane Proteome Array (MPA) platform. The MPA comprises 5,300 different human membrane protein clones, each overexpressed in live cells from expression plasmids that are individually transfected in separate wells of a 384-well plate. The entire library of plasmids is arrayed in duplicate in a matrix format and transfected into HEK293T cells, followed by incubation for 36 h to allow protein expression. Before specificity testing, optimal antibody concentrations for screening were determined by using cells expressing positive (membrane-tethered Protein A) and negative (mock-transfected) binding controls, followed by flow cytometric detection with an Alexa Fluor-conjugated secondary antibody (Jackson ImmunoResearch Laboratories). Cell specific binding was confirmed first. Based on the assay setup results, VH-Fc protein (20 μg/ml) was added to the MPA. Binding across the protein library was measured on an iQue3 (Ann Arbor, MI) using the same fluorescently labeled secondary antibody. To ensure data validity, each array plate contained positive (Fc-binding; MSLN protein) and negative (empty vector) controls. Identified targets were confirmed in a second flow cytometric experiment by using serial dilutions of the test antibody. The identity of each target was also confirmed by sequencing.

### Crystallization, data collection and refinement

MSLN complex with VH 3C9 was prepared by injecting the protein mixture to Superdex 75 column (GE Healthcare, Chicago IL), and concentrated to 12 mg/mL. To allow multi-wavelength anomalous diffraction analysis, selenomethionine (Se-Met) labeled VH 3C9 was also prepared by expressing the protein in E.Coli BL21(DE3) cells, using a minimum media with Se-Met and supplemental amino acids to reduce Se-Met toxicity (L-isoleucine, L-leucine, L-lysine, L-phenylalanine, L-threonine, and L-valine) prior to the induction [31]. The Se-Met incorporation to VH 3C9 was confirmed by mass spectrometry, 15122.52 Da (theoretical mass, 15123.23 Da), by comparing the mass without Se-Met labeling, 15028.68 Da (theoretical mass 15029.41 Da), using a LC–TOF mass spectrometry (Bruker Daltonics, Billerica, MA). Data were collected at the 12-2 SSRL (wavelength 0.9795 Å). Diffraction data were processed, integrated and scaled using XDS. The structure was solved by single-wavelength diffraction analysis. Model building and refinement were performed using Coot and Phenix [32, 33] (Table 1).The complex structure will be deposited to the Protein Data Bank.

### Animal model generation in trials

Animal trials were completed in Abrev Inc (both in vivo ADC efficacy and distribution). Briefly, Nude mice (18-22 g, female, Qls03-0102) were purchased from Qing Long Shan Animal Center (Nanjing, China). The nude mice were kept following Laboratory Animal center’s standard operational procedures. 4-6 week old nude mice were injected via s.c. with 5.0 × 10^5 AsPC-1-luciferase cells. Seven days later, the tumor size was about 200 mm3 and was be visible. Antibodies were then injected intraperitoneally (i.p.) as described below.

### Spatial distribution of VH-Fc 3C9 in vivo

Antibodies were labeled with far infrared dye YF750 SE (US EVERBRIGHT INC, YS0056) (named as Nbs-YF750). Five mice (M01-05) were injected intraperitoneally (i.p) with 200 μg (~10 mg/kg, mice) purified Abs-YF750. Tumor fluorescence was thus measured accordingly. Images were observed at Ex:740 nm/Em:780 nm by NightOWL LB 983 (Berthold, Germany) at the indicated time point. To cofirm the antibody enriched in the tumor, the tumor was dissected at 3 days post antibody infusion (dpi) and 14 dpi (endpoint), respectively. To pinpoint the location of the tumor cells, luciferase substrate was injected via i.p. to measure the signal of luciferase from AsPC-1-luciferase cells. Images were analyzed using Indigo imaging software Ver. A 01.19.01.

## List of Supplementary Materials

Materials and Methods

Fig S1 to S6 for multiple supplementary figures.

Table S1.

## Acknowledgments

We would like to thank the members of the Center for Antibody Therapeutics in University of Pittsburgh for their helpful discussions. We thank Andrew Hinck for the use of the Biacore X100 instrument.

## Author contributions

Z.S. and D.S.D. designed project. Z.S. identified and characterized antibodies, designed CAR-T, DDCs and in vivo assay. X.C. produced recombinant MSLN proteins for crystallization study. C.A. performed UPLC for DAR determination and cell cytotoxicity assays. C.C. produced the VH-Fc protein for assay. D.J. and W.L. studied the modeling of epitope of antibody. M.G., G.L, R.I., and G.C did crystallization. T.I., M.G., and R.I. conducted SPR experiments and SEC experiments to verify the binding site. G.C. determined crystal structure. Z.S. wrote the first draft of the article. C.A. revised the draft. Z.S., J.W.M., and D.S.D. discussed the results and further revised the manuscript. All authors contributed to the final manuscript.

## Data and materials availability

All data are available in the main text or the supplementary materials. The PDB file will be subject to submission after a peer reviewed journal accepts our manuscript.

---

**Figure S1.**
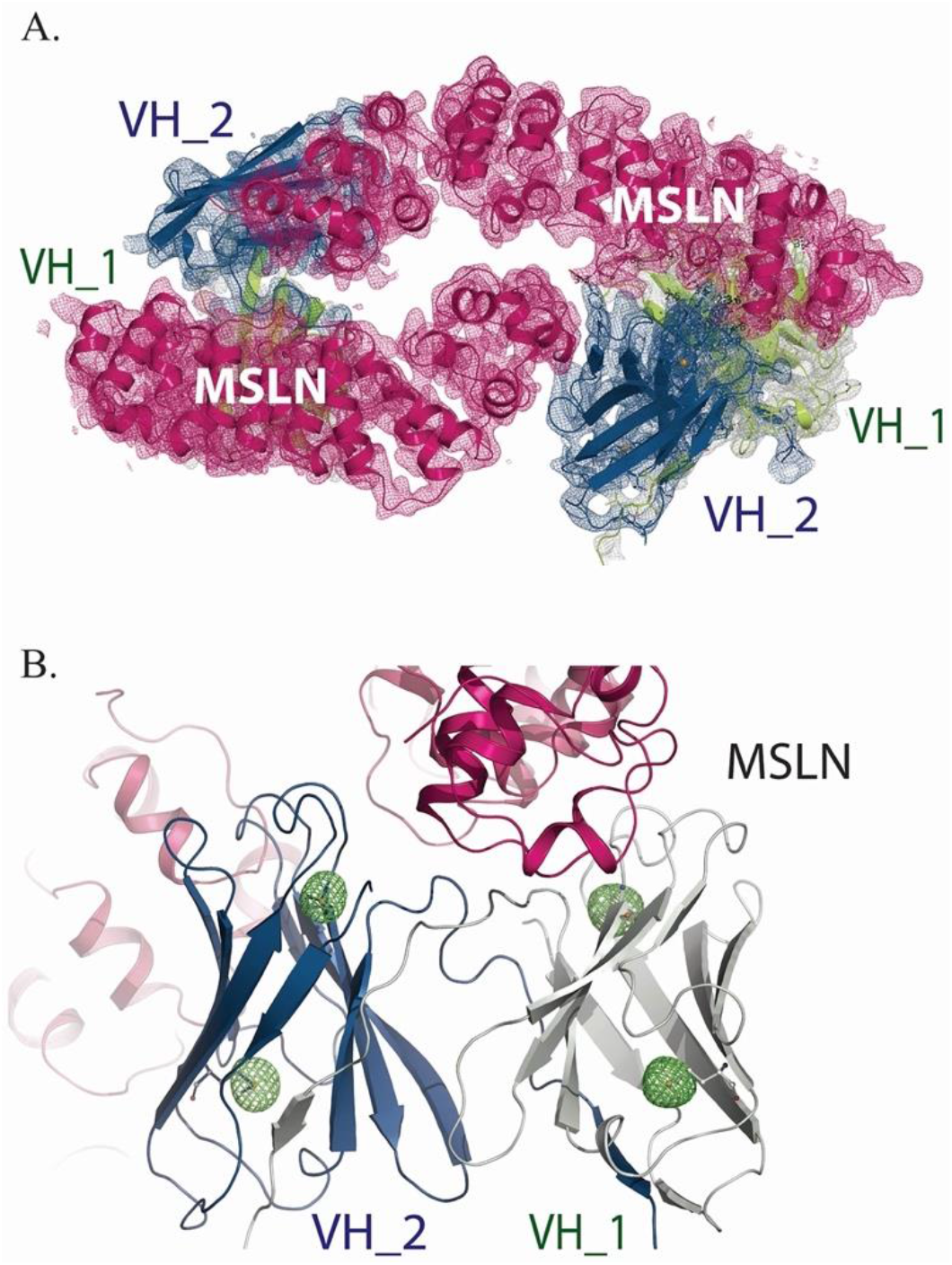
(A) Electron density (2f0) map of MSLN, VH_1, and VH_2 in a unit cell and (B) the positions and the density of seleno-methionines in VH_1 and VH_2.

**Figure S2.**
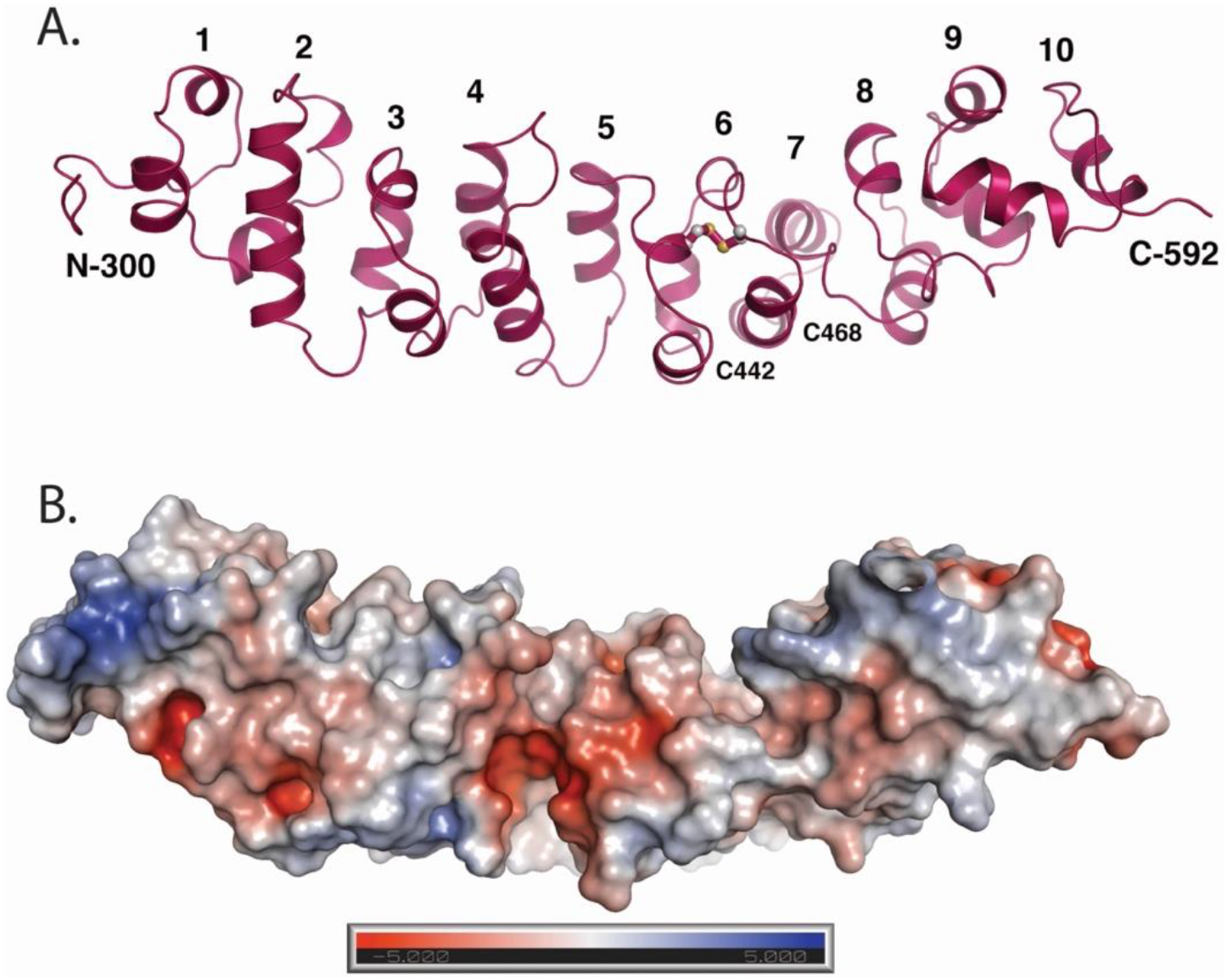
Structural characteristics of MSLN: (A) helical numbers, and (B) electrostatic surface potential of MSLN calculated by PyMol from −5 kT/e (red) to + 5 kT/e (blue).

**Figure S3.**
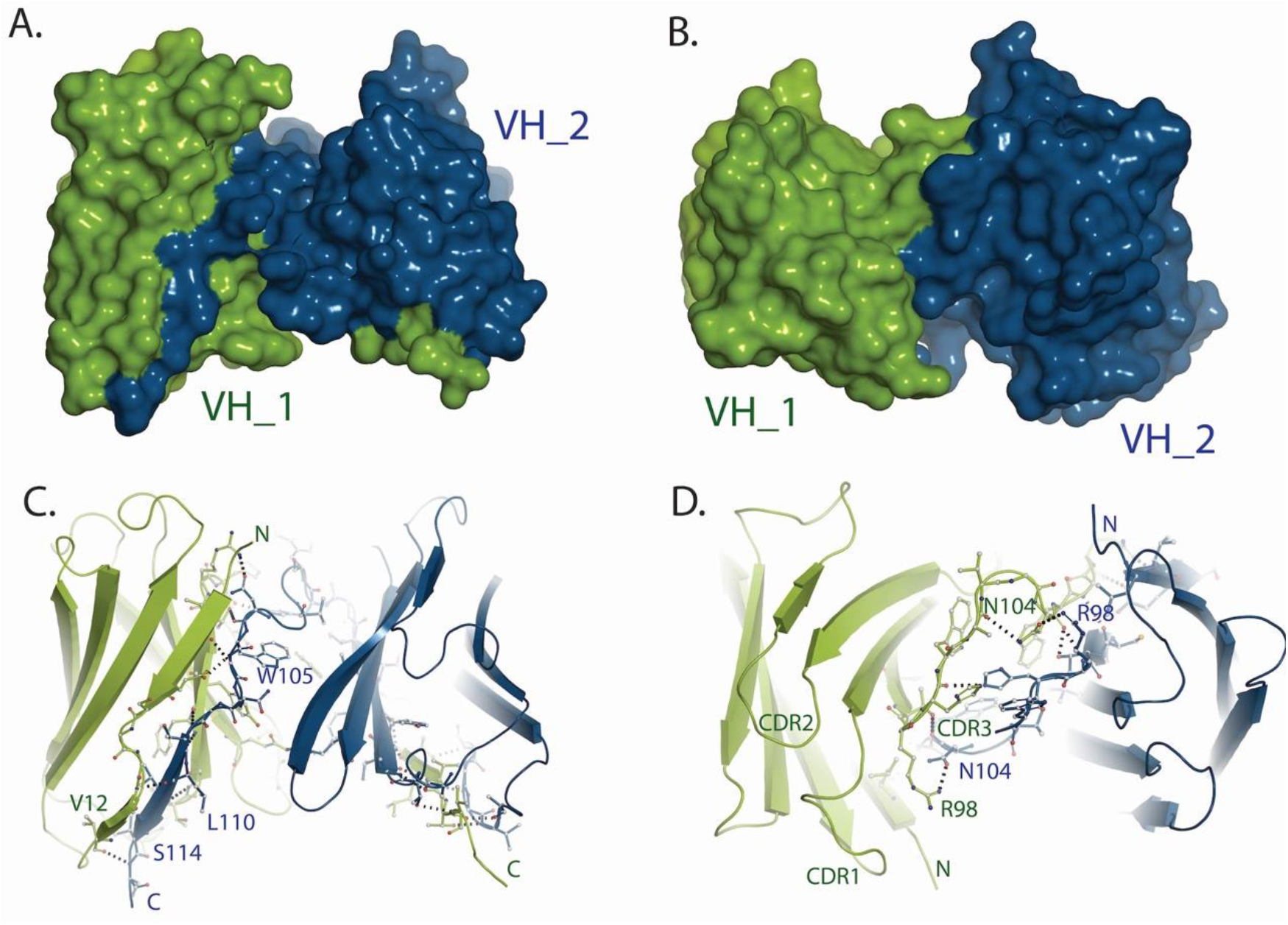
Structural characteristics of the two molecules of VH 3C9, VH_1 (green) and VH_2 (blue): (A) surface presentations of the two molecules, (B) that of the 90-degree rotated view, (C) Side chain interactions of the N-terminal strand in VH_1 and the swapped C-terminal strand in VH_2, and (D) Side chain interactions of the CDR3 regions in the two molecules that interface to each other.

**Figure S4.**
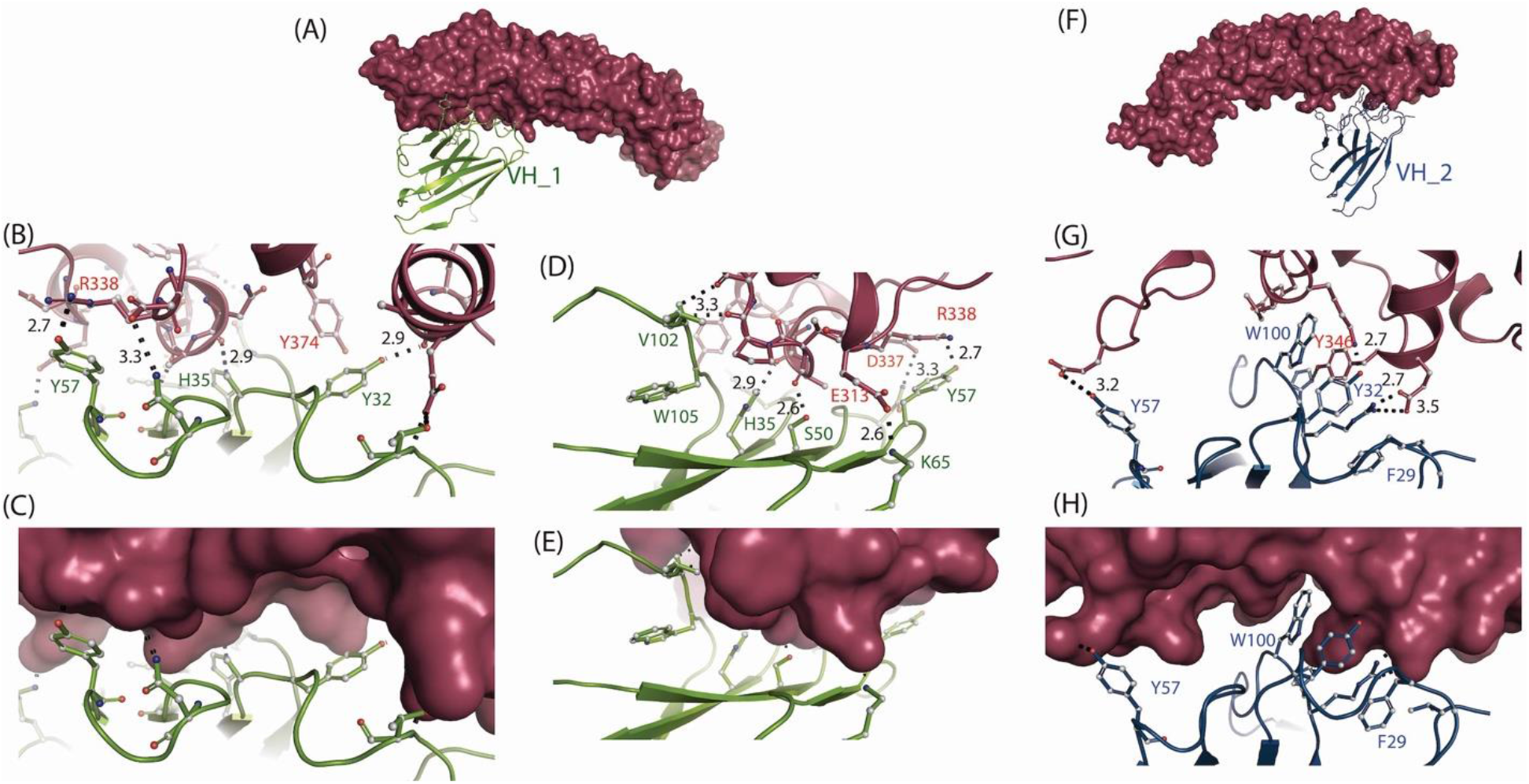
Interactions of MSLN and (A-E) VH_1 and (F-H) VH_2: (A) showing relative orientation of VH_1 against MSLN; (B, C) interaction of the side chains of VH_1 and with surface of MSLN; (C, E) approximately 90 degree oriented view for (B, C) respectively; (F) showing relative orientation of VH_2 against MSLN; (G, H) interaction of the side chains of VH_2 and with surface of MSLN. MSLN, VH_1, and VH_2 are shown by purple, green, and blue colors, respectively.

**Figure S5:**
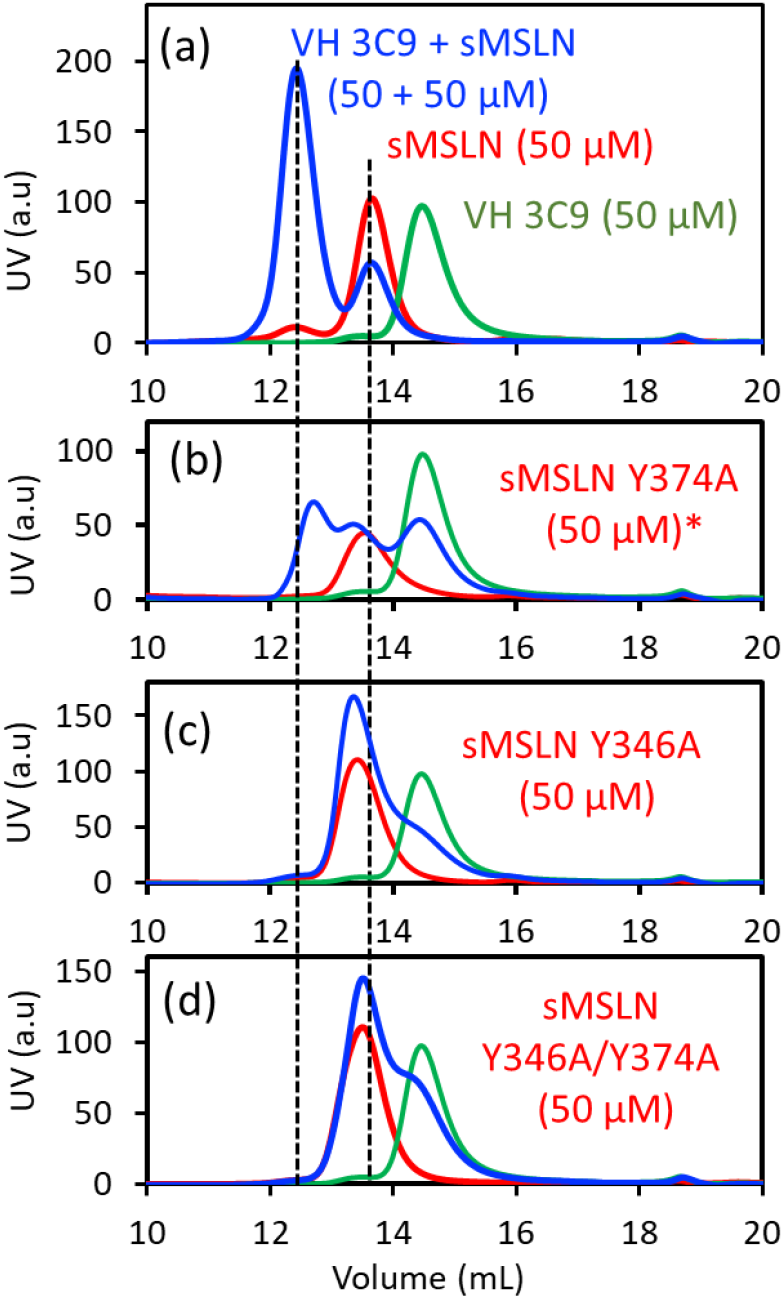
Confirmation of the VH 3C9 binding site on MSLN. Elution profiles of sMSLN alone (red), VH 3C9 alone (green) and the mixture (blue), using (a) wt sMSLN, (b) Y374A sMLSN, (c) Y346A sMSLN and (d) Y346A/Y374A sMSLN. We have purchased a clone of a short MSLN (sMSLN, residues 296 to 415, with N-terminal His-tag and TEV cleavage site, and the C-terminal avi tag) from ATUM, and expressed in *E. coli* BL21 cells. In panel (a), since approximately a half of sMSLN does not interact with VH 3C9 when the equal amount is mixed, sMLSN:VH3C9 binding stoichiometry is estimated to be 1:2. In panel (b), sMSLN Y374A shows binding to VH3C9 weaker than that of WT. In panel (c, d), no binding is observed in Y346A or Y346A/Y374A.

**Figure S6.**
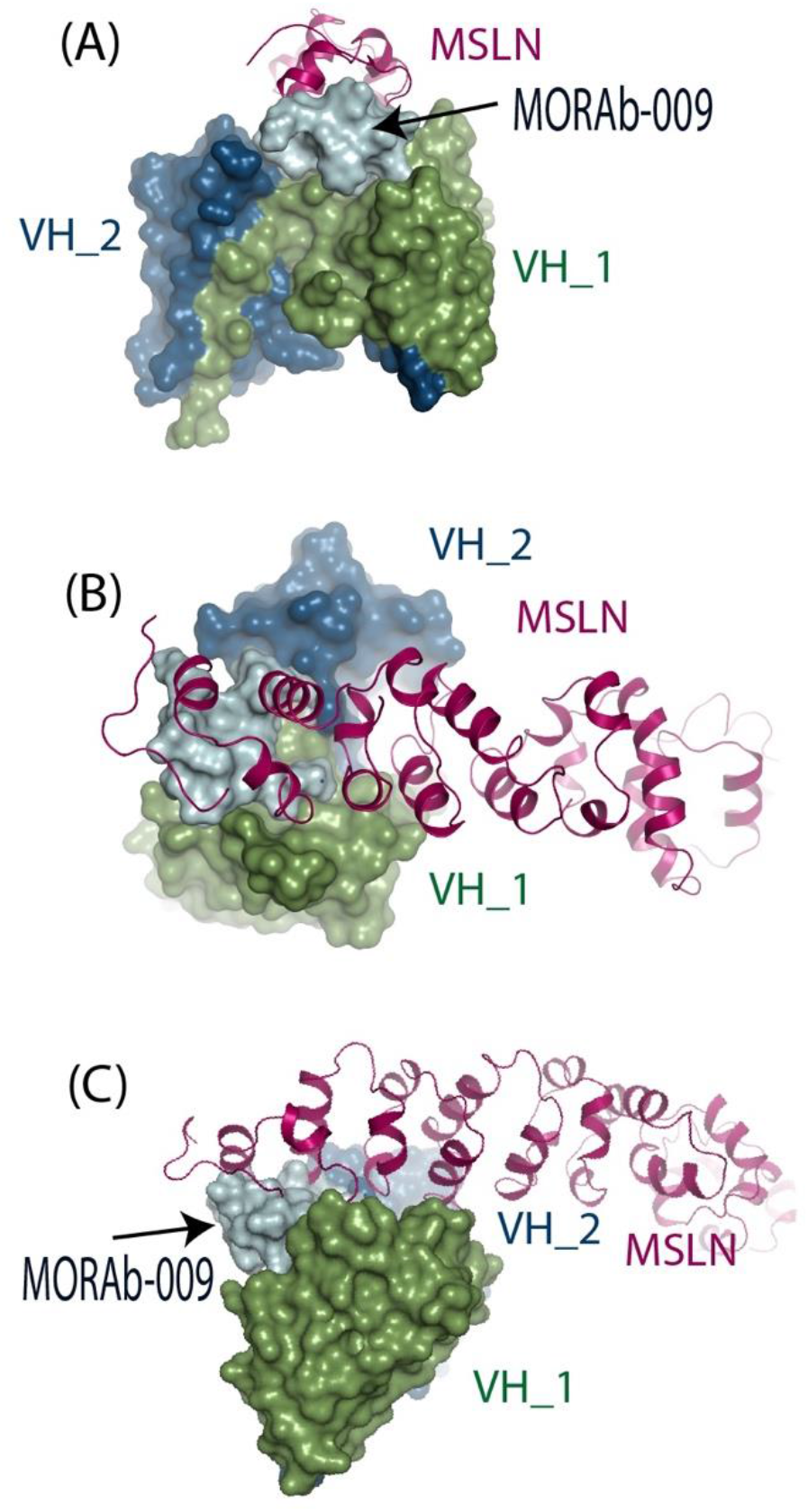
Amatuximab Fab (MORAb-009) interaction site on the N-terminal of MSLN (ice, PDB 4F3F) and VH_1 (green) and VH_2 (blue) extensively interacting with MSLN (purple), shown from three different orientations, A, B, and C.

**Table S1.**
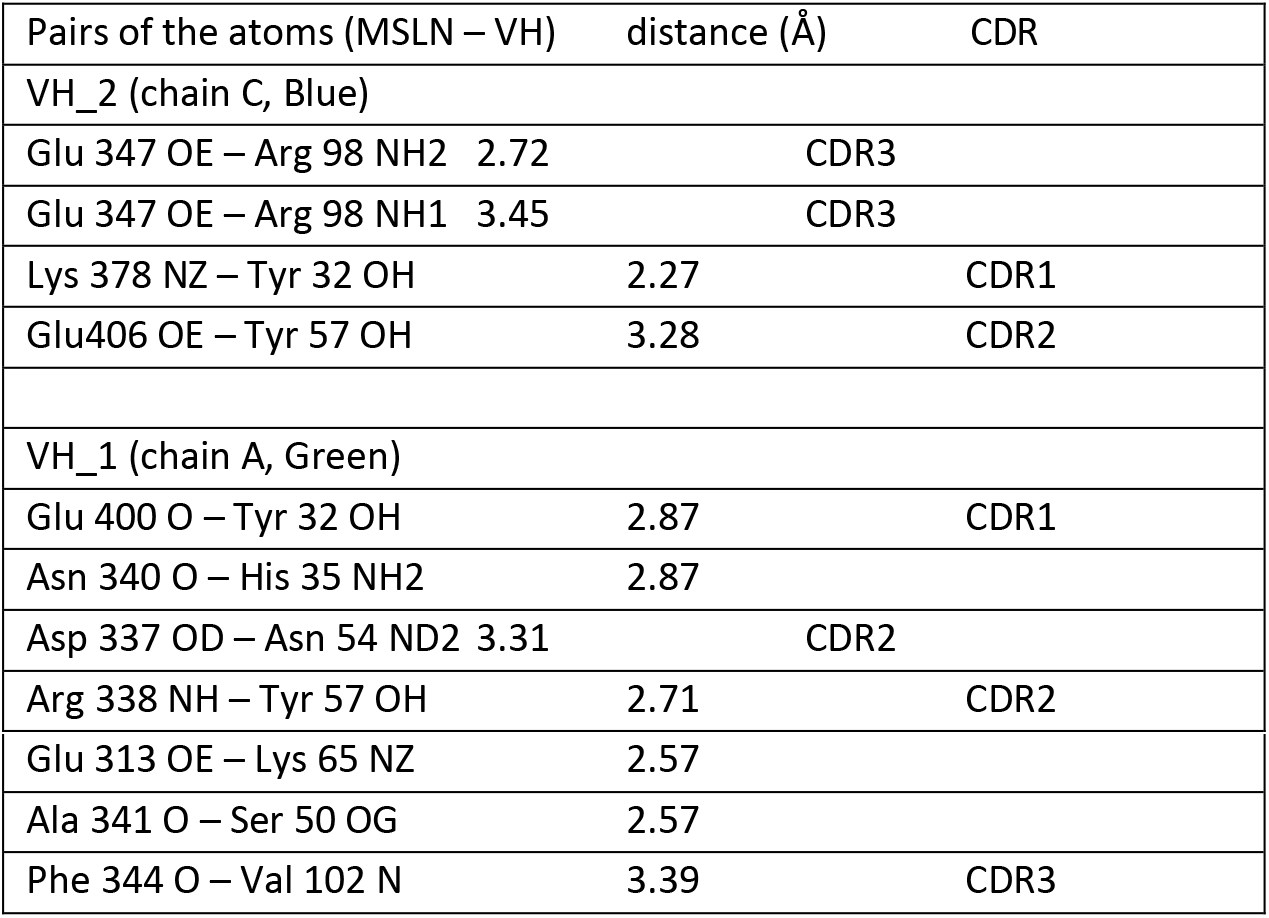
Hydrogen bonds in the MSLN – VH interaction sites.

